# Brain Amyloid and the Transition to Dementia in Down Syndrome

**DOI:** 10.1101/2020.08.19.257790

**Authors:** David B. Keator, Eric Doran, Lisa Taylor, Michael J. Phelan, Christy Hom, Katherine Tseung, Theo G.M. van Erp, Steven G. Potkin, Adam M. Brickman, Diana H. Rosas, Michael A. Yassa, Wayne Silverman, Ira T. Lott

## Abstract

**INTRODUCTION:** Down syndrome (DS) is associated with elevated risk for Alzheimer’s disease (AD) due to beta amyloid (Aβ) lifelong accumulation. We hypothesized that the spatial distribution of brain Aβ predicts future dementia conversion in individuals with DS.

**METHODS:** We acquired ^18^F-Florbetapir PET scans from 19 nondemented individuals with DS at baseline and monitored them for four years, with five individuals transitioning to dementia. Machine learning classification determined features on ^18^F-Florbetapir standardized uptake value ratio (SUVR) maps that predicted transition.

**RESULTS:** In addition to “AD signature” regions including the inferior parietal cortex, temporal lobes, and the cingulum, we found that Aβ cortical binding in the prefrontal and superior frontal cortices distinguished subjects who transitioned to dementia. Classification did well in predicting transitioners.

**DISCUSSION:** Our study suggests that specific regional profiles of brain amyloid in older adults with DS may predict cognitive decline and are informative in evaluating the risk for dementia.

**Highlights:**

- Regional [18F]-Florbetapir PET predicts future transition to dementia in Downs Syndrome.
- Increased amyloid in prefrontal, inferior parietal, superior frontal, rostral middle frontal, and posterior cingulate cortices detect transitioiners, with prefrontal and superior frontal being best overall.
- Amyloid PET-based classification able to discriminate between transitioners and non-transitioners.

## 1. INTRODUCTION

Individuals with Down syndrome (DS) have a high age-related prevalence of Alzheimer’s disease (AD) and life-long accumulation of brain amyloid (Aβ) in part due to the triplication of amyloid precursor protein (APP) on chromosome 21 [1]. Early identification of those at highest risk for early onset dementia is paramount for intervention trials in DS since potential disease modification is less effective after symptoms of cognitive decline are observable [2]. Current approaches for detecting brain Aβ include Positron Emission Tomography (PET) [3,4]. One important issue is whether amyloid-PET alone could be predictive of dementia transition in DS. As Aβ accumulates in the brain across the lifespan in DS, it may be possible to identify regional Aβ distributions that predict future conversion to dementia.

In this study, a small sample of cognitively stable (non-demented) participants with DS were followed clinically for four years after acquiring a baseline [^18^F]-Florbetapir PET scan. During the four-year follow up, a subset progressed to dementia. Here we report on differential amyloid accumulation as a function of transition time and classification results of predicting those who transitioned, using the baseline PET scan. Regions of particular interest of amyloid uptake were the frontal regions, middle and inferior temporal cortices, and the inferior parietal cortex, as these areas have been linked to executive functioning, visuospatial processing, and memory and are consistent with late Braak and Braak stages [5]. We hypothesized these regions would show preferential uptake during dementia transition.

## 2. METHODS

### 2.1 Participant Characteristics

Nineteen cognitively stable adult participants with DS were evaluated at baseline and after nine, 18, 27, and 48 months. Informed consent was obtained according to IRB protocols for persons with intellectual disabilities. During the period of follow-up, five participants progressed (age=50.4+/-4.3yrs.; sex=2 Male, 3 Female; average transition time from PET scan=1.9±1.3yrs.) to dementia based on clinical evaluations; fourteen remained cognitively stable (age=52.1+/- 5.7yrs.; sex=10 male, 4 female). The groups did not differ in mean age (t(17)=0.70;p<0.49 two-tailed) or sex (p<0.30 Fisher’s). Baseline PET scans were acquired along with longitudinal neuropsychological assessments (supplement S3).

### 2.2 Diagnosis of the Transition to Dementia

Dementia was diagnosed in accordance with ICD-10 and DSM-IV-TR criteria as outlined by Sheehan et al. [6]. Transition classification followed comprehensive baseline and longitudinal assessments including history, neurological examination, and consideration of previous studies in the medical record. Transition to dementia was decided at a consensus conference, blinded to the PET scan and neuropsychological results. Participants with confounding conditions (e.g. sensory deficit, untreated thyroid dysfunction, and major depression) were excluded. Details regarding transition symptoms of individual participants are given in the supplemental material (supplement S1).

### 2.3 Image Acquisition

[^18^F]-Florbetapir PET scans were acquired at the University of California, Irvine Neuroscience Imaging Center using the High Resolution Research Tomograph (HRRT) [7]. Image acquisition followed the Alzheimer’s Disease Neuroimaging Initiative (ADNI) [8] protocol. PET reconstructions were performed using the 3D ordinary Poisson ordered subset expectation maximization (3D OP-OSEM) algorithm with scatter and attenuation corrections [9]. Structural T1-weighted MPRAGE scans were acquired on a 3-Tesla Philips Achieva scanner (sagittal orientation, TR/TE=6.8/3.2ms, flip angle=9°, NEX=1, field of view=27cm^2^, voxel resolution=0.94×0.94×1.20mm, matrix size=288×288×170, SENSE acceleration factor=2).

### 2.4 Image Processing

The PET frames were realigned, averaged, and co-registered with their respective MRI scans. MRI segmentations were computed with the FreeSurfer (FS6; RRID:SCR_001847). Regions of interest (ROI) were extracted in the native MRI space from the FS6 Desikan/Killiany atlas [10] segmentations. The PET counts were converted to standardized uptake value ratio (SUVR) units using the cerebellum cortex reference region. Correction for partial volume effects was performed using PETSurfer [11]. Voxel-weighted ROI averages for inferior parietal, entorhinal, lateral occipital, anterior/posterior cingulate, inferior/middle/superior temporal, prefrontal, superior frontal, rostral-middle frontal, medial/lateral orbito-frontal, precuneus, dorsal striatum, hippocampus, and whole-brain were computed for each subject. To understand whether regional amyloid burden was predictive of transition, we modeled the logit of the probability of transition as a linear function of average amyloid burden for each ROI after removing the linear effects of region volume and age.

## 3. RESULTS

In the following sections, we describe analyses designed to evaluate: (1) the relationship between transition time to clinical dementia and regional amyloid burden; (2) the effect size of differences between transitioned and non-transitioned participants; (3) our ability to predict who would transition to frank dementia.

### 3.1 Brain Amyloid and Risk of Clinical Conversion

Cox regression models were built with the R survival package to evaluate the risk of conversion given the variable transition times relative to the PET scan acquisitions (RRID:SCR_001905). Separate regression models were evaluated for each ROI after removing the linear effects of region volume on ROI SUVR average and including age at the time of PET scan as a covariate. The results are shown in Table 1 where regions are ordered by magnitude of risk. The Cohen’s D effect sizes of the group differences for each ROI is included.

**Table 1.** Cox regression analysis results with and without partial volume correction (PVC) by region of interest. Table shows the z-score from Cox regression, uncorrected p-values (P), adjusted (Hommel method) p-values (P Adj), and Cohen’s D effect size of transitioned vs. non-transitioned participants.

| Regions of Interest | PVC |  |  |  | Without PVC |  |  |  |
| --- | --- | --- | --- | --- | --- | --- | --- | --- |
|  | Z | P | P Adj | Cohen's D | Z | P | P Adj | Cohen's D |
| BrainAverage | 2.53 | 0.006 | 0.044 | 1.87 | 2.51 | 0.006 | 0.041 | 1.19 |
| SuperiorFrontal | 2.53 | 0.006 | 0.044 | 1.86 | 2.50 | 0.006 | 0.041 | 1.85 |
| Prefrontal | 2.24 | 0.012 | 0.056 | 1.81 | 2.39 | 0.009 | 0.047 | 1.80 |
| MiddleTemporal | 2.45 | 0.007 | 0.047 | 1.74 | 2.00 | 0.026 | 0.077 | 2.18 |
| PosteriorCingulate | 2.25 | 0.012 | 0.056 | 1.67 | 2.44 | 0.007 | 0.045 | 2.18 |
| RostralMiddleFrontal | 2.20 | 0.014 | 0.056 | 1.66 | 2.37 | 0.009 | 0.047 | 1.65 |
| InferiorParietal | 2.64 | 0.004 | 0.037 | 1.60 | 2.71 | 0.003 | 0.030 | 1.77 |
| SuperiorTemporal | 2.20 | 0.014 | 0.056 | 1.55 | 2.32 | 0.010 | 0.051 | 1.43 |
| Precuneus | 2.62 | 0.004 | 0.040 | 1.52 | 2.61 | 0.004 | 0.035 | 1.65 |
| LateralOrbitoFrontal | 1.92 | 0.027 | 0.074 | 1.29 | 2.11 | 0.017 | 0.063 | 1.90 |
| MedialOrbitoFrontal | 1.98 | 0.024 | 0.072 | 1.18 | 2.26 | 0.012 | 0.052 | 1.24 |
| AnteriorCingulate | 1.97 | 0.024 | 0.073 | 1.13 | 2.13 | 0.017 | 0.063 | 1.29 |
| DorsalStriatum | 1.83 | 0.033 | 0.074 | 1.10 | 1.86 | 0.031 | 0.094 | 0.95 |
| InferiorTemporal | 1.65 | 0.049 | 0.098 | 0.89 | 2.65 | 0.004 | 0.032 | 1.62 |
| Hippocampus | 0.18 | 0.431 | 0.431 | 0.13 | 0.66 | 0.254 | 0.259 | 0.35 |
| EntorhinalCortex | -1.39 | 0.083 | 0.083 | -0.73 | -0.46 | 0.323 | 0.323 | -0.23 |
| LateralOccipital | -1.93 | 0.027 | 0.054 | -1.32 | 0.65 | 0.259 | 0.259 | 0.09 |

### 3.2 Amyloid Classification of Future Clinical Transition

Given the robust findings from the regression analyses, we were interested in evaluating the performance of a classification algorithm in detecting future dementia transitions using amyloid PET data prior to conversion. Because of the small sample size, we used an independent dataset of eleven participants with DS from the Alzheimer’s Biomarkers Consortium-Down Syndrome (ABC-DS) with an initial consensus-based diagnosis of mild cognitive impairment-Down Syndrome (MCI-DS) [12], six of which transitioned (range: 1.0-1.6 yrs. after PET scan) to a diagnosis of dementia (age=53.1+/-4.3yrs.;1 woman, 10 men). This allowed us to test the logistic regression classifiers trained with the primary dataset. Using the PET scan data from the independent test set, prior to transition and after removing the linear effects of region volume and age, we evaluated whether the trained classifiers could discriminate between those who transitioned to dementia and those who did not (Table 2). We found, similar to the Cox regression analysis, that amyloid burden in prefrontal, inferior parietal, superior frontal, rostral middle frontal, and posterior cingulate were among the most sensitive regions for detecting who would eventually transition. Broadly looking across the metrics, we found prefrontal and superior frontal cortices to be the best overall regions, with high sensitivity, balanced accuracy, area under the receiver operating characteristic (auc) curve, and reasonably good specificity (0.80).

**Table 2:**
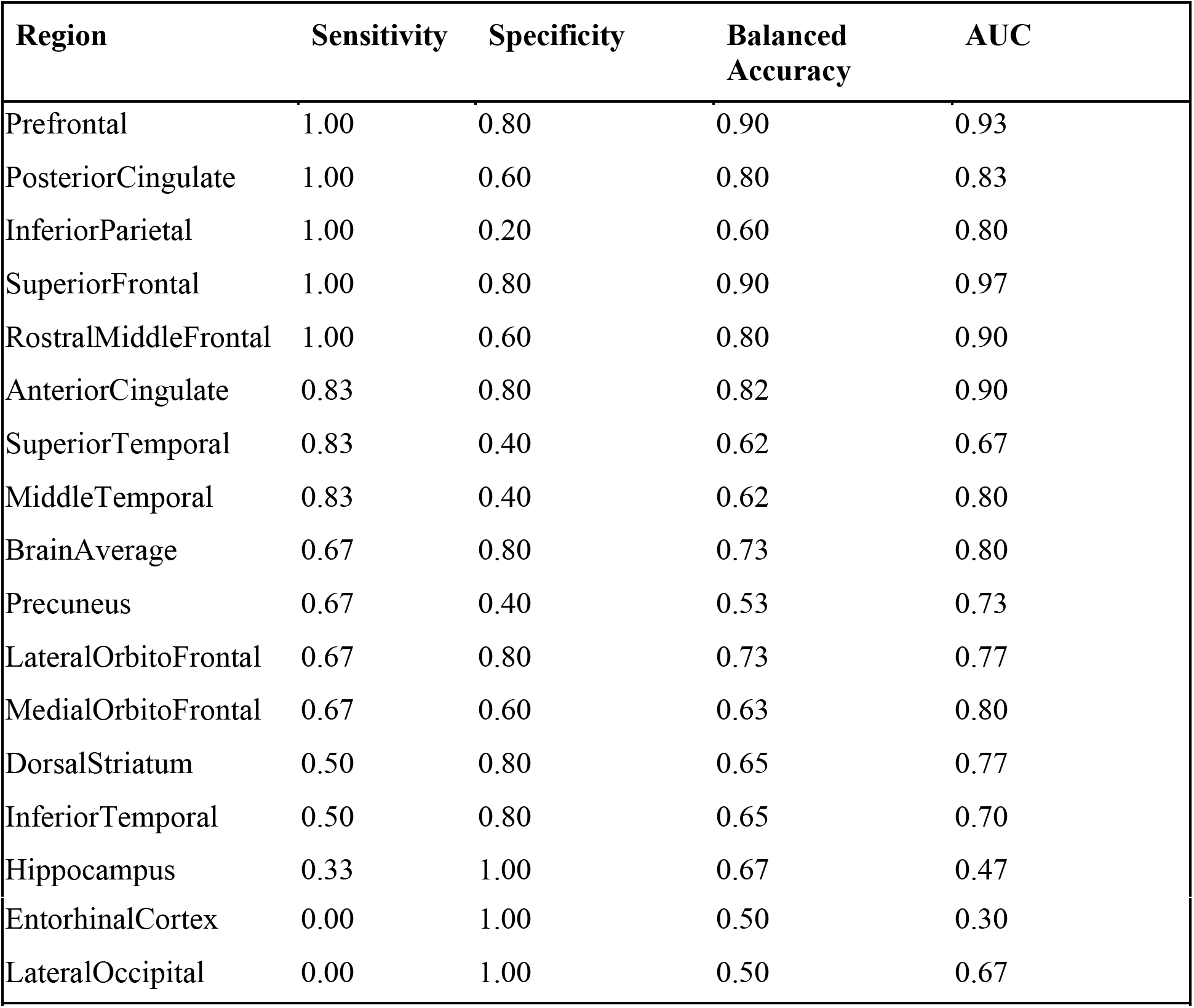
Logistic regression classifier results tested on an independent sample of participants with MCI (N=11) prior to conversion to dementia (N=6). Metrics shown include specificity, sensitivity, accuracy balanced for number of participants in each test group, and area under the ROC curve (auc).

## 4. DISCUSSION

This retrospective case-control study evaluated a cohort of nineteen participants with DS, five of whom transitioned to dementia during the course of follow-up. The primary goal was to evaluate the regional distribution of amyloid prior to clinical transition and understand whether amyloid alone predicts future transition. Exploiting an advantage of logistic regression with retrospective study designs [13], these analyses described the predictive potential for classifying cases based on regional amyloid distribution in scans prior to transition. We found that high amyloid burden in several regions, namely the prefrontal, superior and rostral middle frontal and posterior cingulate provided excellent prediction of dementia transition. These regions are broadly associated with executive function, working memory, and attentional focus which have been implicated in dementia progression in both DS [14] and neurotypical populations [15].

In the neurotypical population, the biomarker utility of increased cortical uptake as a result of amyloid binding on PET scans has been of interest in preclinical AD. However, it is not clear that the uptake data reliably predicts those who will subsequently transition to dementia [3]. Various working groups in nuclear medicine and AD have developed criteria for using amyloid PET in the diagnosis of patients with persistent or unexplained mild cognitive impairment in the neurotypical population but the diagnostic and predictive status of these measures remains uncertain [16]. Other groups have found the rate of β-amyloid accumulation in the brains of individuals with DS differ according to the pre-existing amyloid burden [17,18] and the presence of cortical Pittsburgh compound B (PiB)-binding, another PET amyloid ligand, as a function of age which has been confirmed in post-mortem studies [19]. Our study suggests regional profiles of brain amyloid in older adults with DS may predict cognitive decline and are informative in evaluating the risk for dementia. In a related study, we found increased amyloid in specific regions to be associated with MCI-DS, supporting the use of regional amyloid in tracking dementia progression in DS [12]. Our current study reinforces these observations, suggesting a role for regional amyloid measurements in a composite risk score for predicting dementia progression in individuals with DS. Pending replication, regional amyloid is likely a useful quantitative measure to include in a composite risk score for dementia transition in DS.

## Acknowledgments and Funding

This work was supported in part by NICHD 065160 (Lott), R01 AG053555 (May), and P50 AG16573 (May). Independent dataset for classifier testing was provided by the Alzheimer’s Disease in Down Syndrome (ADDS) component of the Alzheimer’s Biomarkers Consortium –Down Syndrome (ABC-DS), a longitudinal study of Alzheimer Disease biomarkers in adults with Down syndrome is supported by grants from the National Institute on Aging (NIA) (U01AG051412-01 Schupf, Lott, Silverman) and Eunice Kennedy Shriver National Institute of Child Health and Human Development (NICHD). We would like to acknowledge Sharon Krinsky-McHale, Ph.D. from the New York State Institute for Basic Research in Developmental Disabilities and Margaret Pulsifer, Ph.D. from Massachusetts General Hospital for their input on the interpretation of neuropsychological assessments and results of this study.

## Notes

### Competing Interest Statement

The authors have declared no competing interest.

