## Supplementary Materials for "Brain Amyloid and the Transition to Dementia in Down Syndrome"

### Supplementary Material

### S1. Description of participants who transitioned to dementia

Participant 1

Age 47 years

Female

10 visits between age 35 and 46 years

This woman has a 12 month history of cognitive decline. She frequently loses track of her personal possessions having been previously meticulous in caring for them. She is less oriented to familiar surroundings in the community and now gets lost at her favorite movie house. She is less oriented to the calendar and now will inaccurately recall the correct day of the week, month, year and season. She is having difficulty remembering the birthdays of close family members and the names of friends. She has lost the ability to accurately keep score during bowling games. Her productivity at her workshop has declined. It takes her longer to produce speech and it is less intelligible and more simplified. On a daily basis, verbal prompting and some assistance is now required for her to carry out her dressing and toileting. There have been a number of urinary accidents. She is now easily angered and experiences anxiety when separated from her caregiver. She has had recent episodes of choking while eating and is eating too quickly. Her gait has become more hesitant and she often requests support when approaching door thresholds, curbs or stairs.

Participant 2

Age 57 years

Female

7 visits between ages 52 and 57 years

There is a 2 year history of progressive difficulties with memory and overall abilities. She gets confused frequently in attempts to carry out commands. She is now having difficulty remembering recent events. She often requires assistance in carrying out daily activities for which she was previously independent. She has lost her previous ability to dress according to the weather. She consistently loses personal possessions. She is less oriented to familiar surroundings and has difficulty finding her way within her home. She has become very emotional and has frequent crying spells. She has developed aggressive behaviors, particularly when she doesn't get her own way. She is often anxious in social settings and will withdraw from group activities. She has lost motivation for activities she previously enjoyed such as television and coloring. Even though she had articulation problems throughout her life, her speech has become more difficult for others to understand.  Her gait has slowed and she hesitates frequently when crossing a threshold or transition in flooring pattern, often requesting that her hand be held. She is no longer able to copy her name and other simple drawings. She has developed myoclonic seizures.

Participant 3

Age 59 years

Male

9 visits between ages 49 and 61 years

He has a 1 ½ year history of a progressive decline in his memory and functioning. As a result, he tends to ask the same question repeatedly. Having previously been independent for scheduling most his daily activities, he now relies on others and becomes confused in the timing of his activities. He is no longer oriented to the calendar and is forgetting his own birthday. He is having difficulty remembering the details of recent events and the names of people familiar to him. He has stopped reading the daily newspaper and no longer follows his favorite sports teams. He is less engaged in social conversations and activities at his workshop, and often isolates himself from others. He has shown a recent loss of inhibition and will now make inappropriate comments to others. While living semi-independently, he has shown increased reliance on a part-time caretaker for activities that he had mastered previously, including his hygiene and showering andnow puts on dirty clothes from the day before. There is occasional bowel and bladder incontinence. His gait has become slower and less coordinated resulting in severalfalls. There were 2 pathological reflexes on neurological examination.

Participant 4

Age 58 years

Female

9 visits between ages 53 and 58 years

This woman has experienced a progressive decline in her memory level of functioning over the past 3 years. She is requiring more prompts to carry out her activities of daily living and now requires hand-over-hand assistance for bathing and dressing. Previously meticulous about her appearance, she now refuses to shower or change clothes. She has become intolerant of frustration --often lashing out and hitting people. She has become destructive of property within the house. She has become anxious and upset when her caregiver of 8 years is not nearby. She now needs 24-

hour a day supervision.  Her moods are easily changeable and will quickly go from laughter to tears. She now believes her parents, who have been deceased for over a decade, are visiting her. She now has urinary incontinence. There is some reversal in her sleep cycle, with her being awake most of the night and sleeping during the day. As a result, she is sleepy in her day program and refuses to participate. Her gait has become slower and less coordinated and she now requires physical assistance with stairs and curbs. There was a single pathological reflex on neurological exam.

Participant 5

Age 52 years

Male

6 visits between 48 and 52 years

This previously high-functioning man began to show a progressive decline beginning 1.5 years ago. He is now forgetting steps in his routine activities of daily living such as remembering to rinse the shampoo from his hair when showering. As a result of his confusion during daily activities, he had to transition to a higher level of care. He also wanders away and gets lost. He now has difficulty with 2- and 3-step commands and understanding complex sentences. He now recalls less details about past events.. His productivity in his workshop declined to the point that he had to leave his workshop and he attended a day program instead. He neglects his appearance, for which he was previously fastidious, and now wears torn or dirty clothing. His speech is becoming less articulate and his sentences are less complex. He often requires support while walking and has lost his ability to tandem walk during the most recent neurological exam. There have been several severe coughing and choking episodes at mealtime.

### S2. Voxel-based Effect Sizes of Increased Amyloid in Transitioners

To estimate effect sizes of increased amyloid in participants who transitioned, a voxel-based linear regression was performed in SPM12 using ANCOVA scaling, scan age, and transition time. Maps of t-statistics contrasting increases in amyloid in the transitioned group were converted to estimated effect-size maps (Figure S2) using the f-modeling approach to empirical Bayes estimation [43]. Regions with large effect sizes are labeled in Figure S1 to show correspondence with the ROI Cox regression results. Reviewing the t-map contrast using a probability threshold of $p<0.001$ uncorrected ($t(14)=3.78$) demonstrated significant increases in the superior and middle frontal, medial orbit-frontal, superior temporal, precuneus, posterior cingulate, and inferior parietal.

Generally, the voxel-based analyses were consistent with the ROI results, with large effect sizes (> 2.0) in the superior frontal, rostral middle frontal, inferior parietal, superior/middle temporal, and posterior cingulate regions ($p<0.001$unc).

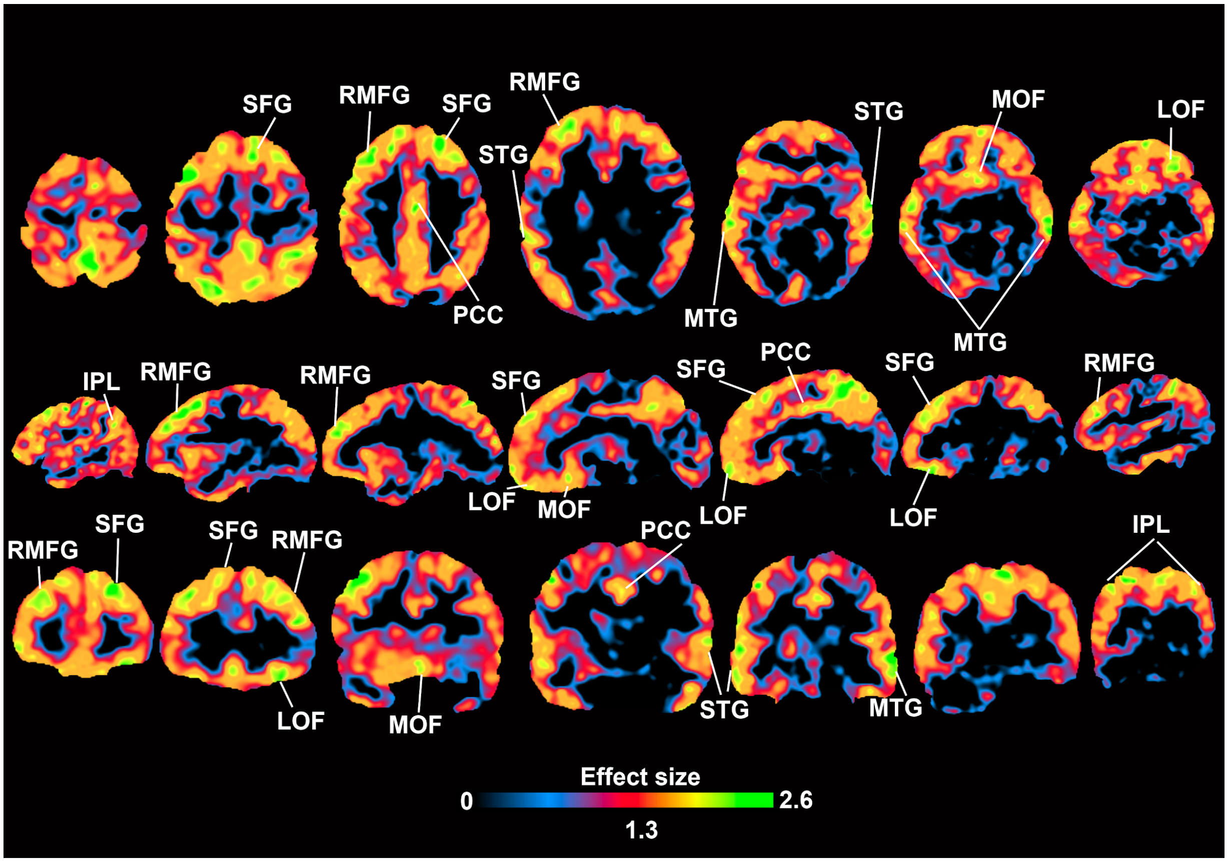


Figure S1: Voxelwise effect size estimates of transitioned group compared to non-transitioned group. Positive effect sizes indicate areas where the transitioned group had increased amyloid relative to the non-transitioned group. Regions are labeled with abbreviations: IPL, inferior parietal lobe; LOF, lateral orbitofrontal; MOF, medial orbitofrontal; MTG, middle temporal gyrus; RMFG, rostral middle frontal gyrus; PCC, posterior cingulate cortex; SFG, superior frontal gyrus; STG, superior temporal gyrus.

### S3. Longitudinal Neuropsychological Outcomes and Baseline Amyloid

The following longitudinal assessments were collected on participants: Severe Impairment Battery (SIB), Rapid Assessment for Developmental Disabilities (RADD), Fuld Object Memory Evaluation (FULD), and an informant/caregiver questionnaire, Dementia Questionnaire for people with Learning (DLD, formerly the Dementia Questionnaire for Persons with Mental Retardation (DMR)) Sum of Cognitive Scores (SCS) and DLDDMR Sum of Social Scores (SOS). The participants who transitioned to dementia during the longitudinal observation period showed progressive problems with memory, gait, ability to perform activities of daily living, and decreased productivity in their workshop and community settings. The mean rates of decline and the differences between the transitioned and non-transitioned groups on the neuropsychological measures are shown in Table S2 and are consistent with a diagnosis of dementia in the group that transitioned. For the RADD, SIB, and FULD measures, a negative rate of change indicates a worsening of performance. For the DMR SCS and DMR SOS measures, a positive rate of change indicates a worsening of performance. The difference between the average rates of decline on the SIB, for example, in the transitioned group was estimated to be 5.05±1.25 points per year faster than in the non-transition group. Differences on the other measures can be interpreted in a similar fashion.

| Diagnosis | N | RADD | SIB | FULD | DMR-SCS | DMR-SOS |
|---|---|---|---|---|---|---|
| **Non-Transition (NT)** | 14 | 0.62 ± 0.59 | 1.23 ± 0.46 | 1.66 ± 1.54 | -0.51 ± 0.28 | -0.27 ± 0.52 |
| **Transitioned (T)** | 5 | -2.64 ± 0.69 | -3.82 ± 1.16 | -2.95 ± 1.12 | 2.18 ± 1.33 | 2.23 ± 0.87 |
| **Difference (NT-T)** | | 3.26 ± 0.91 | 5.05 ± 1.25 | 4.61 ± 1.91 | -2.69 ± 1.36 | -2.51 ± 1.02 |

S2: Mean rate of decline in neuropsychological performance by diagnostic group.

To evaluate whether higher brain amyloid burden at baseline for each ROI was associated with a steeper rate of change in neuropsychological performance, we performed a series of regression analyses across the entire group of subjects, irrespective of transition status (n=19) after removing the linear effects of MRI volume. Overall, the SIB was the only measure that had a significant relationship with regional amyloid at the uncorrected $p<0.05$ level (Table S3).

Interestingly, unlike the risk analysis, whole brain average amyloid did not significantly relate to longitudinal SIB decline nor decline in the other assessments.

| | PVC | | | Without PVC | | |
|---|---|---|---|---|---|---|
| **Region of Interest** | **T Value** | **P Value** | **P Adj** | **T Value** | **P Value** | **P Adj** |
| **PosteriorCingulate** | -2.93 | 0.011 | 0.165 | -2.99 | 0.010 | 0.138 |
| **Precuneus** | -2.55 | 0.023 | 0.277 | -2.66 | 0.019 | 0.206 |
| **DorsalStriatum** | -2.54 | 0.023 | 0.281 | -2.64 | 0.019 | 0.209 |
| **BrainAverage** | -2.47 | 0.027 | 0.306 | -1.76 | 0.101 | 0.404 |
| **LateralOrbitoFrontal** | -2.18 | 0.046 | 0.405 | -3.08 | 0.008 | 0.122 |
| **InferiorTemporal** | -1.98 | 0.067 | 0.405 | -2.17 | 0.048 | 0.325 |
| **AnteriorCingulate** | -1.94 | 0.072 | 0.405 | -2.09 | 0.055 | 0.331 |
| **MedialOrbitoFrontal** | -1.91 | 0.077 | 0.405 | -2.11 | 0.053 | 0.325 |
| **SuperiorTemporal** | -1.60 | 0.131 | 0.405 | -1.69 | 0.114 | 0.423 |
| **MiddleTemporal** | -1.47 | 0.164 | 0.405 | -1.98 | 0.068 | 0.371 |
| **Prefrontal** | -1.46 | 0.165 | 0.405 | -1.43 | 0.173 | 0.476 |
| **InferiorParietal** | -1.31 | 0.210 | 0.405 | -1.90 | 0.078 | 0.392 |
| **Hippocampus** | -0.93 | 0.371 | 0.405 | -1.04 | 0.317 | 0.635 |
| **RostralMiddleFrontal** | -0.88 | 0.396 | 0.405 | -1.39 | 0.186 | 0.476 |
| **SuperiorFrontal** | -0.88 | 0.396 | 0.405 | -1.18 | 0.257 | 0.515 |

| | | | | | | |
|---|---|---|---|---|---|---|
| **EntorhinalCortex** | -0.86 | 0.405 | 0.405 | -2.38 | 0.032 | 0.257 |
| **LateralOccipital** | 0.99 | 0.340 | 0.405 | -0.37 | 0.716 | 0.716 |

Table S3: Relationship between baseline amyloid and longitudinal rate of change in SIB assessment. Negative t-values indicate that higher baseline amyloid load is associated with a larger decline in SIB performance.

### S4. Neuropsychological Rates of Change Comparison

The mean rates of decline and the differences between the transitioned and non-transitioned groups on the neuropsychological measures are shown in Table S4. A MANOVA was used to compare the rates of change in cognitive ability with age, gender, handedness, transition status, and the interaction between gender and clinical transition status. There was a main effect of transition status (Pillai's trace=0.81, F=7.84, df=5,9, p<0.004) and a main effect of gender (Pillai's trace=0.78, F=6.53, df=5,9. p<0.008) on the rates of decline on the neuropsychological tests. Transition status was significantly related to RADD slope (F=14.40, p<0.002), SIB slope (F=33.87, p<5.97e-05), DMR SCS slope (F=7.27, p<0.02), and DMR SOS slope (F=10.41, p<0.007). Further, there was a significant gender x transition status interaction on RADD slope (F=4.98, p<0.04), SIB slope (F=12.48, p<0.004), FULD slope (F=5.86, p<0.03), DMR SCS slope (F=5.19, p<0.04), and DMR SOS slope (F=8.60, p<0.01). In light of the gender x transition status interactions, adjusted mean differences by gender were computed and shown in Table S4. The unadjusted average rate of decline on the SIB, for example, in the transitioned group was estimated to be 5.05+/-1.25 points per year faster than in the non-transition group. However, this difference was significantly modified by the sex of the participant. The adjusted average difference in rate among females (F) was estimated to be 8.21+/-1.20 points per year, but only 1.46+/-1.30 points per year among males. Mean differences for other scales in Table S4 may be interpreted using the same approach and were significant for females ($p<0.001^{**}$ and $p<0.005^{*}$) but not for males.

| Diagnosis | N | RADD | SIB | FULD | DMR-SCS | DMR-SOS |
|---|---|---|---|---|---|---|
| Non-Transition (NT) | 14 | 0.62±0.59 | 1.23±0.46 | 1.66±1.54 | -0.51±0.28 | -0.27±0.52 |
| Transitioned (T) | 5 | -2.64±0.69 | -3.82±1.16 | -2.95±1.12 | 2.18±1.33 | 2.23±0.87 |
| Unadjusted Diff (NT-T) | | 3.26±0.91 | 5.05±1.25 | 4.61±1.91 | -2.69±1.36 | -2.51±1.02 |
| Adjusted Diff (NT-T) | F<br>M | $^{**}$6.32±1.45<br>1.20±1.58 | $^{**}$8.21±1.20<br>1.46±1.30 | $^{*}$12.29±3.53<br>-0.56±3.83 | $^{*}$-4.49±1.24<br>-0.01±1.34 | $^{*}$-5.79±1.12<br>0.24±1.21 |

Table S4: Mean (± standard error) longitudinal rates of change per year on neuropsychological measures. The transitioned group had worsening performance over time across all tests.
